# A mouse model of metastasis-induced breast cancer cachexia with non-cachectic controls bearing isogenic primary tumors

**DOI:** 10.64898/2026.10.28.762598

**Authors:** Yanshan Liang, Young-Yon Kwon, Marie Sabatier, Mario Palma, Jessalyn M. Ubellacker, Sheng Tony Hui

## Abstract

**Background:** Patients with advanced cancer frequently develop cachexia, an involuntary weight-loss condition that correlates strongly with metastatic burden. Despite this clinical association, metastasis is rarely considered in preclinical research of cancer cachexia, which is in part due to the lack of rigorously established animal models to investigate metastasis-associated cancer cachexia.

**Methods:** To establish an animal model of metastasis-associated cancer cachexia, here we focused on syngeneic mouse models of breast cancer. Using the metastatic 4T1 murine breast cancer cell line and its isogenic non-metastatic 67NR cell line, we generated primary tumors via orthotopic injection into the mammary fat pad and surgically resected the primary tumor to allow sufficient time for spontaneous metastatic outgrowth, or generated experimental lung metastases by direct intravenous injection into the tail vein. Across these tumor-bearing groups, we systematically characterized the metastatic burden and cachectic phenotypes of the mice, including body weight, food intake, body composition, muscle mass, grip strength, locomotor activity, energy expenditure, and respiratory exchange ratio. To isolate the contribution of reduced food intake to the cachectic phenotypes, we pair-fed orthotopic 4T1 tumor-bearing mice (which were non-cachectic) to mice with 4T1 lung metastases (which were cachectic). We further measured muscle atrophy gene expression and profiled circulating cytokines and metabolic hormones for the mouse groups.

**Results:** Body weight loss occurred exclusively in mice bearing lung metastases from 4T1 tumors, with no weight loss observed in mice bearing either primary 4T1 tumors or 67NR lung tumors, thus presenting a model of cancer cachexia that is driven by 4T1 lung metastasis. Comprehensive cachexia characterization demonstrated that the 4T1 metastasis model recapitulates all key features of cachexia, manifesting as reduced food intake (or anorexia), depletion of adipose tissues, loss of lean mass including muscle mass loss, and reduced grip strength. Pair-feeding non-cachectic primary 4T1 tumor-bearing mice to 4T1 metastatic mice led to the same losses in body weight, fat, and lean mass as in cachectic mice, but no change in grip strength, indicating that anorexia is the primary driver of tissue wasting whereas muscle weakness arises through a food intake-independent mechanism. Moreover, we identified distinct circulating cytokines and metabolic hormones associated with the metastasis-driven cachexia.

**Conclusions:** Our results establish a metastasis-induced breast cancer cachexia model with a tightly matched control group, providing a platform for the field to identify metastasis-derived cachectic mediators and pathways.

## 1 Introduction

Cancer cachexia is a debilitating metabolic syndrome characterized by involuntary weight loss that compromises patient quality of life, treatment tolerance, and overall survival^1,2^. Prevalent in advanced cancer, cachexia frequently occurs in parallel with metastatic dissemination^3–5^. Despite this strong association, metastasis has been largely ignored in cancer cachexia studies. One contributor to this situation is the lack of well-established in vivo metastasis models that capture the co-existence of both metastasis and cachexia. The most widely used animal models of cancer cachexia include a colon carcinoma 26 (C26) mouse model^6^ and the Lewis lung cancer (LLC) mouse model^7^, neither of which readily undergo spontaneous metastasis in vivo. Even in commonly used GEMMs of cancer cachexia, including those for pancreatic ductal adenocarcinoma (PDAC)^8^ and non-small cell lung cancer (NSCLC)^9^, whether there is metastasis when cachexia takes place is usually not reported.

There are a limited number of investigations using metastasis models of cancer cachexia^10–17^. Among them are transplant models using metastatic breast cancer cell lines. Waning et al. inoculated a human breast cancer cell line (MDA-MB-231) to the left cardiac ventricle to induce bone metastases^18^. Regan et al. also obtained bone metastases by injecting the 4T1 mouse breast cancer cell line to the right tibia^19^. In both studies, muscle weakness and wasting were reported. Two additional studies used the spontaneous metastasis models in which orthotopically implanted 4T1 tumor cells are surgically resected to permit outgrowth of spontaneous metastasis (primarily lung), and reported body weight loss and tissue wasting^11,17^. While valuable results have been gained from using the models in these studies, a rigorously controlled, comprehensive characterization of cachexia is lacking, preventing a wide adoption of a metastatic model of cancer cachexia. For example, none of the studies had measured energy expenditure, which is critical for evaluating energy balance of the condition. Moreover, none controlled for metastatic tumor burden in affected organs (e.g., the lungs), which itself can impair organ function and contribute to cachectic phenotypes. Thus, there is a need in the field for a rigorously controlled and systematically characterized animal model of metastatic cancer cachexia.

In this study, building on previous studies ^11,17,19^, we focused on transplant models using the 4T1 murine breast cancer line. We investigated both the surgical resection (of orthotopic primary tumors) as well as an intravenous (IV) injection approach as a tool to induce experimental metastasis. To control for tumor burden, we included in our experiments the 67NR cell line, which is an isogenic, non-metastatic counterpart to the 4T1 cell line. Through systematic characterization of cachexia as well as tumor burden for mice bearing either primary or metastatic tumors, we identified mice with IV injection of 4T1 cells as a robust model of metastasis-induced cancer cachexia. For this model, we characterized all main aspects of cachexia, comprising body weight, adipose and lean mass, muscle mass, and physical strength, as well as energy balance parameters including food intake and energy expenditure. We further determined the effects of reduced food intake on cachectic phenotypes and performed inflammatory and metabolic hormonal profiling in the model. Altogether, our results provided a robustly established mouse model for studying metastasis-induced cancer cachexia.

## 2 Materials and Methods

### 2.1 Cell culture

The 4T1 breast cancer cell line was purchased from ATCC, and the 67NR cell line was a kind gift from Dr. Robert A. Weinberg (MIT). All cells were maintained in incubators at 37 °C and 5% CO_2_, using RPMI 1640 Medium (Corning) with 10% fetal bovine serum (FBS, R&D Systems) and 1% penicillin/streptomycin (P/S, Hyclone).

### 2.2 Generation of luciferase-tagged cell lines

4T1 cells and 67NR cells expressing iRFP720 and luciferase were generated by lentiviral transduction. HEK293T cells were co-transfected with 5 µg of pHIV-iRFP720-E2A-Luc (Addgene plasmid #104587), 5 µg of psPAX2 (Addgene plasmid #12260), and 0.5 µg of pMD2.G (Addgene plasmid #12259) packaging plasmids using Lipofectamine 3000 according to the manufacturer’s instructions. Viral supernatants were collected 48 and 72 h after transfection and passed through a 0.45 µm filter. For transduction, 4T1 and 67NR cells were incubated with the filtered viral supernatant supplemented with 8 µg/mL Polybrene (Sigma-Aldrich, H9268). The transduction procedure was performed twice. Following transduction, iRFP720-positive cells were isolated by fluorescence-activated cell sorting (FACS). Luciferase expression was confirmed by incubating the cells with D-luciferin and measuring bioluminescence using an IVIS imaging system.

### 2.3 Animals

BALB/c female mice were purchased from Charles River Laboratories and were housed (4 per cage) under regular light-dark cycles of 12 h with ad lib access to water and food (PicoLab 5053, LabDiet). Prior to tumor inoculation, mice were single-housed and acclimated to experimental conditions for 1 week. Tumor injection was conducted when animals were 12-16 weeks old and had reached a stable stage in growth and weight gain. For orthotopic injection, 5 × 10^5^ 4T1 or 67NR cells suspended in 200 µL saline were injected into the mammary fat pad of each mouse. For intravenous injection, 2.5 × 10^4^ 4T1 or 1.5× 10^5^ 67NR cells were injected into the circulation through the tail vein. The much higher number of injected 67NR cells than 4T1 cells was necessary for achieving similar tumor growth at the lungs between the two cell lines. For primary tumor resection, we performed survival surgery experiments. Briefly, mice bearing 0.8-1.0 cm (in length) primary tumors were anesthetized using 3% isoflurane and maintained at 37^◦^C, and underwent primary tumor resection using standard sterile surgical procedures. After the surgery, mice received 0.5 mg/kg buprenorphine-SR for analgesia and were monitored for 5 days. Mice were euthanized upon reaching the experimental endpoint, defined as the day when mice lost approximately 15% of initial body weight. Animal care and experimental procedures were conducted with the approval of the Institutional Animal Care and Use Committees (IACUC) of the Harvard Medical Area.

### 2.4 Food intake and pair feeding

Food intake was measured daily at a consistent time (between 10:00 and 11:00□am) and was conducted as we published previously^20^. Briefly, animals were housed individually on alpha pad bedding with kraft paper nesting material. Each day between 4:00 and 6:00 pm, remaining pellets were weighed and returned to the cage lid, and any food debris in the bedding was sieved out (2 mm mesh), weighed, and discarded. Food surplus was the weight of the remaining pellets plus the debris. Daily food intake was the amount of food provided minus the food surplus over two consecutive days.

For the pair-feeding experiment, mice were assigned to three body weight-matched groups at baseline: mice bearing 4T1 primary tumors fed ad libitum, mice bearing 4T1 lung metastases fed ad libitum, and mice bearing 4T1 primary tumors that were pair-fed. Beginning on day 10 post-tumor injection, daily food intake of the lung metastasis group was recorded, and the pair-fed group received a matched amount (the preceding day’s average intake). To mimic the nocturnal feeding pattern of ad libitum-fed mice, the allotted food was delivered automatically in three equal portions (7:00pm, 11:00pm, and 3:00am) using an automatic fish feeder (Petbank). Cages were checked daily to confirm that the entire allotment had been consumed.

### 2.5 Body composition assessment

Whole-body composition (fat and lean mass) was assessed for each animal on day 0 and endpoint using an EchoMRI-100 (EchoMRI LLC). Mice were individually placed in a specialized plastic tube for mild conscious restraint and scanned for approximately 5 minutes. Following measurement, mice were returned to their home cages.

### 2.6 Metabolic cage analyses

Mice were individually housed at 22 °C on alpha pad bedding (6“ × 10”, ScottPharm solution) and subjected to indirect calorimetry right after tumor inoculation under a 12 h light-dark cycle using the Promethion Metabolic Cage system (Sable Systems, USA). During this period, spontaneous activity, oxygen consumption (VO□), and carbon dioxide production (VCO□) were continuously measured to calculate energy expenditure (EE) and respiratory exchange ratio (RER).

### 2.7 Grip strength

Grip strength was assessed using a Grip Strength Meter (Bioseb); mice were allowed to grasp a metal grid with all four limbs and were gently pulled backward until release, with the peak force recorded.

### 2.8 RNA extraction and RT-qPCR

Fresh gastrocnemius, diaphragm and heart samples were isolated from mice and immediately snap-frozen using liquid nitrogen. Total RNA was extracted using the Trizol reagent (Invitrogen, #15596026) from the 20-30 mg of gastrocnemius and TA muscle tissues, following the manufacturer’s protocol. RNA concentration was determined using NanoDrop One (Thermofisher Scientific). One microgram of purified RNA was converted to cDNA using the iScript Reverse Transcription Supermix for RT-qPCR (Bio-Rad, #1708840). The diluted cDNA was used for RT-qPCR using SsoAdvanced Universal SYBR Green Supermix (Bio-Rad, #1725270) with the CFX384 Real-Time PCR detection System (Bio-Rad). The following primer sequences were used to determine the *Trim63*, *Fbxo32*, *Bnip3*, and *Map1lc3b* mRNA expression. *Cyclophilin* was used as a housekeeping gene for normalization.

### 2.9 Plasma metabolic hormones and cytokines

To measure circulating metabolic hormones, blood was collected using K3EDTA-coated collection tubes (Microvette® 500 EDTA K3E, Sarstedt) treated with protease inhibitor cocktails and DPPIV added immediately. Plasma was obtained by centrifugation at 2,000g for 10 min at 4°C. Metabolic hormones in the plasma were quantitated by Eve Technologies (Calgary, Canada) using Mouse/Rat Metabolic Hormone Discovery Assay® Array. Cytokines in the plasma were quantitated by Eve Technologies (Calgary, Canada) using Mouse Cytokine 68-Plex Discovery Assay®.

## 3 Results

### 3.1 Metastatic 4T1 tumors, but not primary tumors, induce body weight loss

A fundamental consideration in establishing a model of metastatic cancer cachexia is whether the primary tumor itself induces cachexia. If so, the model may have limited utility for distinguishing cachectic factors and pathways specifically driven by metastatic disease from those associated with primary tumor growth. To address this question in the case of 4T1 models, we first monitored body weight for mice injected with 4T1 cells at the orthotopic site of the mammary fat pad. These mice did not lose body weight even when the primary tumor was close to 1.0 gram (Fig. 1A). Thus, at the primary site, 4T1 tumors do not cause cachexia.

**Figure 1.**
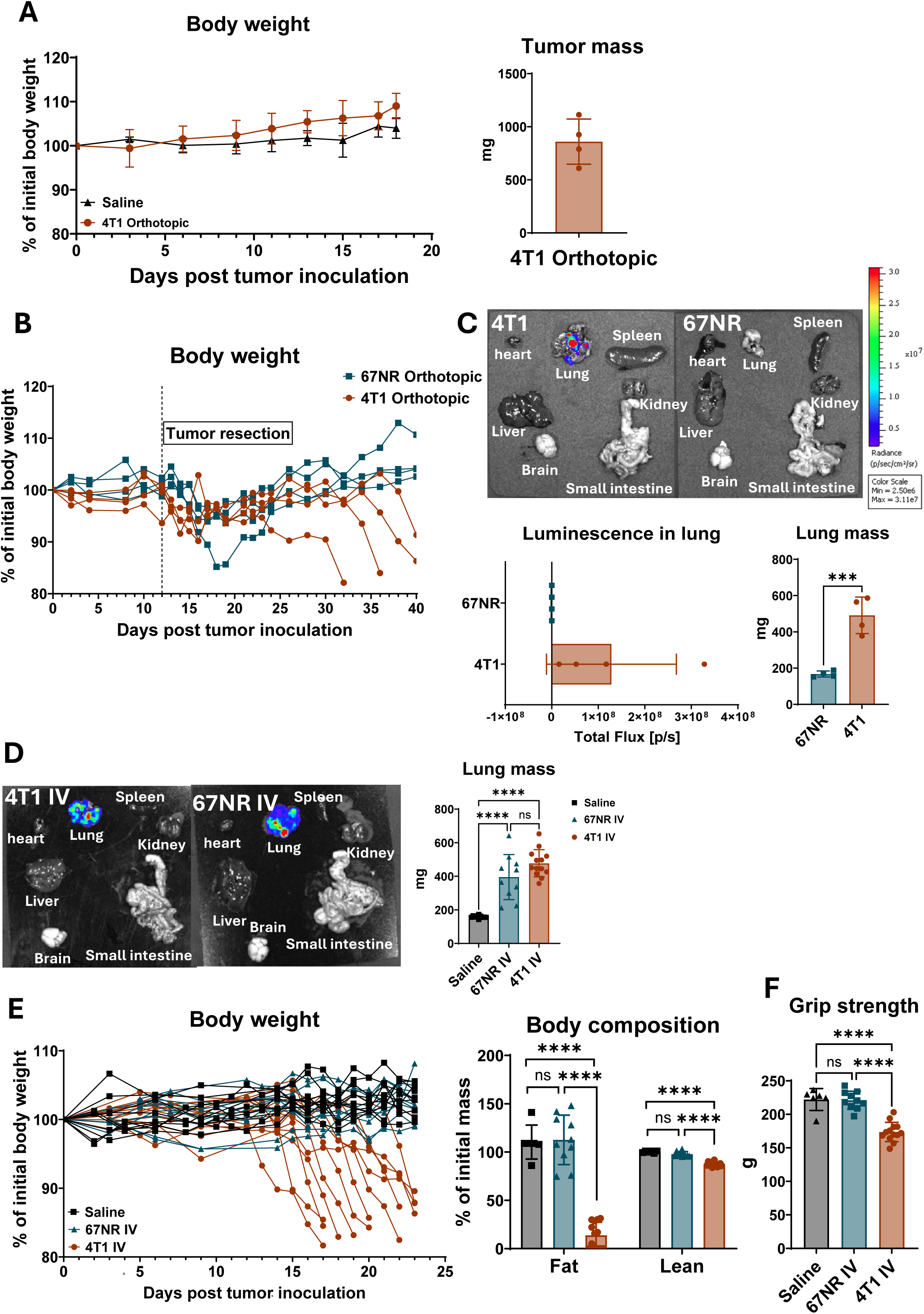
Metastatic 4T1 tumors, but not primary tumors, induce cachexia independently of lung tumor burden. (A) Left panel: Body weight over time in saline-injected non-tumor bearing mice (n=3) or 4T1 orthotopic tumor-bearing mice (n=4); Right panel: Primary tumor mass in 4T1 orthotopic tumor-bearing mice at day 18 post tumor inoculation. (B) Body weight over time in mice bearing orthotopic 67NR (n=4) or 4T1 (n=4) tumors; the primary tumor was resected at day 12 (dashed vertical line) post tumor inoculation (C) Upper panel: representative ex vivo bioluminescence images of harvested organs (heart, lung, spleen, liver, kidney, brain, small intestine) from mice bearing orthotopic 4T1 or 67NR tumors at the experimental endpoint. Primary tumors were resected at day 12 post-tumor inoculation. Lower panels: Quantification of bioluminescence signal (from luciferase-tagged 67NR or 4T1 cells in the tumor-bearing mice) in the lung (lower left), and lung mass at experimental endpoint (lower right). n=4 mice per cohort. (D) Left panel: Representative ex vivo bioluminescence images of harvested organs from mice injected intravenously (IV) with 4T1 or 67NR cells at the experimental endpoint. Right panel: Lung mass at the experimental endpoint of mice injected intravenously with saline (n=10), 67NR cells (n=10), or 4T1 cells (n=13). (E) Left panel: Body weight over time in mice injected intravenously with saline (n=10), 67NR cells (n=10), or 4T1 cells (n=13). Right panel: Body composition at experimental endpoint in mice injected intravenously with saline (n=6), 67NR cells (n=10), or 4T1 cells (n=13). (F) Grip strength in mice injected intravenously with saline (n=6), 67NR cells (n=10), or 4T1 cells (n=13). Data are presented as individual data points with mean ± SD in the bar plots. Statistical significance was determined by one-way ANOVA with Tukey’s multiple comparisons or unpaired two-tailed t-test. *p□<□0.05; **p□<□0.01; ***p□<□0.001; **** p<0.0001; ns, not significant.

Next, to evaluate cachexia in the setting of metastatic disease, we resected orthotopic 4T1 primary tumors 12 days after tumor cell injection, allowing subsequent assessment of mice bearing metastatic disease in the absence of the primary tumor. As a non-metastatic control, mice were orthotopically injected with 67NR cells, which are isogenic to 4T1 cells but do not metastasize. Both cell lines were engineered to express luciferase to allow quantification of tumor burden throughout the body by bioluminescence. We found that 4T1-injected mice exhibited body weight loss while 67NR-injected mice did not, albeit showing a dip immediately after the resection surgery (Fig. 1B). Consistently with their metastatic ability, metastases were detected (in the lungs) in the 4T1-injected mice and no metastases were detected in the 67NR-injected mice (Fig. 1C). Thus, metastatic lung 4T1 tumors, but not the primary tumors, induce body weight loss.

Although the resection-based metastasis model can be used to study cachexia, it has an important limitation for identifying metastasis-specific drivers and mechanisms of cachexia. Primary tumors regrew at the resection site (Fig. S1A), making it difficult to determine whether the observed weight loss was driven by metastatic disease or by the recurrent primary tumor. To account for this limitation, we decided to also use intravenous injection of the 4T1 cell line as a tool for inducing experimental metastasis in the absence of a primary tumor. Similar to the resection-induced metastatic mice, mice injected intravenously with 4T1 tumors exhibited exclusive lung metastasis (Fig. 1D) and significant loss in body weight, fat mass, and lean mass (Fig. 1E). These mice further showed clear reduction in grip strength (Fig. 1F). As primary tumors were absent in these mice, these cachectic phenotypes were solely due to the metastatic tumors, making the IV-injection model more suitable than the resection-based model for elucidating cachectic factors and pathways.

### 3.2 Lung metastatic 4T1 tumors cause cachexia independent of lung tumor burden

The metastatic tumor burden in lungs was large, and one important concern was that the wasting phenotype could be simply due to damaged lung function as a result of large tumor mass in lungs. To test this, we intravenously injected the 67NR cells to mice. Although 67NR cells metastasized poorly from the mammary fat pad, they were able to colonize lungs when delivered directly into the circulation via intravenous tail vein injection (Fig. 1D). In our experiment, we injected to mice such an amount of 67NR cells that produced a comparable lung tumor burden to the 4T1 lung tumor burden (Fig. 1D), and did not observe in these mice any significant body weight loss (Fig. 1E), body composition (Fig. 1E), or grip strength (Fig. 1F). Thus, the cachectic phenotypes in mice with 4T1 lung metastases were not simply due to tumor mass in lungs.

Though 67NR-injected mice did not show body weight loss at the endpoint despite having a compatible lung mass as the 4T1-injected group, with extended observation, they did eventually lose weight, indicating that sufficient pulmonary burden can itself provoke wasting (Fig. S1B). A regression of body weight loss on lung mass shows that 67NR-injected mice began to lose weight only once total lung mass approached 600 mg (Fig. S1B). In contrast, 4T1-injected mice initiated weight loss at a lung mass of approximately 250 mg and had lost around 20% of initial body weight at lung masses of 600 mg (Fig. S1B). This analysis confirms that 4T1-induced cachexia is driven by properties of the tumor cells rather than by the physical burden of tumors in lungs.

### 3.3 Mice bearing primary 4T1 tumors as a strict control group

Although mice bearing either 67NR lung tumors or primary 4T1 tumors could serve as controls for mice with cachexia associated with 4T1 lung metastases, primary 4T1 tumor-bearing mice provide a more appropriate control for identifying tumor-derived cachectic factors because they control for genetic differences between tumor cell lines. Both the group with primary 4T1 tumors and the group with 4T1 lung metastasis received the same 4T1 cells in the same immunocompetent host and differed only in the site at which tumors were established. Comparisons between these two groups isolate the contribution of metastatic tumors from that of primary tumors. Thus, throughout the rest of this study, we used orthotopic 4T1 tumor-bearing mice (hereafter termed ‘4T1 orthotopic group’) as the control group for mice bearing 4T1 lung metastases established by intravenous injection (hereafter termed ‘4T1 IV group’). As confirmation, side-by-side measurements of the two groups showed that the 4T1 IV group began to lose body weight approximately 15 days after intravenous injection and declined progressively until reaching endpoint, whereas the 4T1 orthotopic group maintained weight over the same period (Fig. 2A).

**Figure 2.**
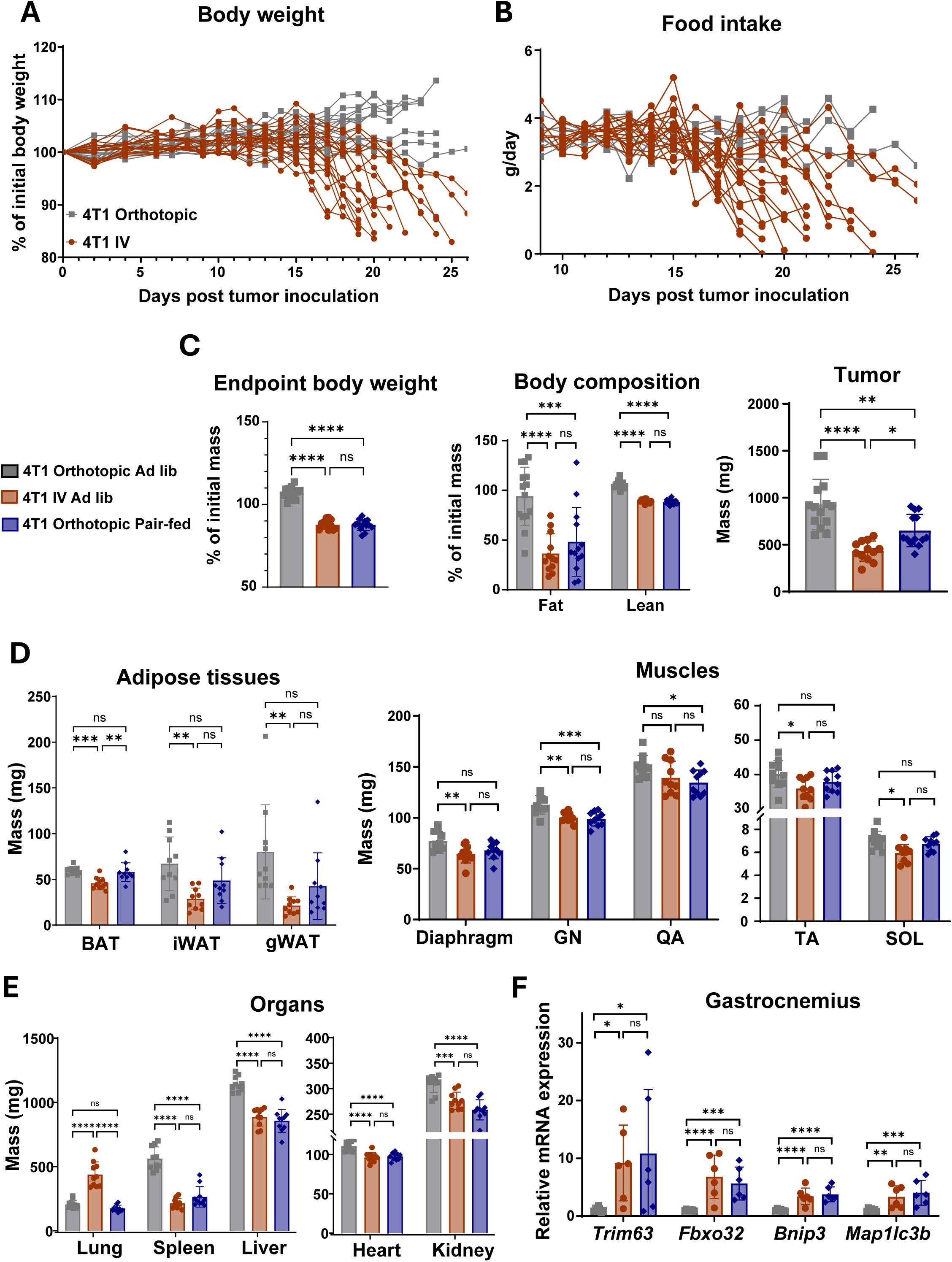
Reduced food intake is the primary driver of body weight loss and tissue wasting in the model of 4T1 metastasis-induced cancer cachexia. (A) Body weight over time in 4T1 IV (n=18) vs. 4T1 orthotopic tumor-bearing mice (n=18). (B) Daily food intake over time in 4T1 IV (n=18) vs. 4T1 orthotopic tumor-bearing mice (n=7). (C) Endpoint body weight, body composition, and tumor mass in 4T1 orthotopic ad libitum mice (n=14), 4T1 IV ad libitum mice (n=12), and 4T1 orthotopic pair-fed mice (n=13). Tumor mass for the 4T1 IV group was obtained by subtracting the lung mass of the pair-fed 4T1 orthotopic littermates from the lung mass of the 4T1 IV mice. (D) Mass of adipose tissues and muscles at endpoint in 4T1 orthotopic ad libitum-fed (n=10), 4T1 IV ad libitum-fed (n=10), and 4T1 orthotopic pair-fed groups (n=10). BAT: brown adipose tissue, iWAT: inguinal white adipose tissue, gWAT: gonadal white adipose tissue, GN: gastrocnemius, QA: quadriceps, TA: tibialis anterior, SOL: soleus. (E) Mass of organs at endpoint in 4T1 orthotopic ad libitum-fed (n=10), 4T1 IV ad libitum-fed (n=10), and 4T1 orthotopic pair-fed groups (n=10). (F) Muscle atrophy and autophagy gene expression in gastrocnemius collected at the experimental endpoint from 4T1 orthotopic ad libitum-fed (n=6), 4T1 IV ad libitum-fed (n=7), and 4T1 orthotopic pair-fed groups (n=6). Data are presented as individual data points with mean ± SD in the bar plots. Statistical significance was determined by one-way ANOVA with Tukey’s multiple comparisons. *p□<□0.05; **p□<□0.01; ***p□<□0.001; **** p<0.0001; ns, not significant.

### 3.4 Reduced food intake is the primary driver for body weight loss and tissue wasting in the model of 4T1 metastasis-induced cancer cachexia

To characterize the model of 4T1 metastasis-induced cancer cachexia, we first measured food intake of mice, revealing a progressive fall in food intake in the 4T1 IV group, mirroring its body weight curve (Fig. 2B). Reduced food intake can be a main driver for key cachectic phenotypes such as body weight loss and tissue wasting^20^. To determine the extent to which cachexia is driven by reduced food intake in our model, we included in our analysis a third group of orthotopic 4T1 tumor-bearing mice that were pair-fed (hereafter referred to as ‘4T1 orthotopic pair-fed group’) to the 4T1 IV group. At endpoint, relative to the 4T1 orthotopic group, the 4T1 IV group and the 4T1 orthotopic pair-fed group had lost the same amount of body weight (Fig. 2C), with similar reduction in both fat and lean mass (Fig. 2C). Importantly, 4T1 orthotopic pair-fed mice had similar tumor mass as 4T1 IV mice, allowing them to control for tumor mass (Fig. 2C).

The same pattern held across individual tissues. Inguinal and gonadal white adipose depots were significantly depleted in the 4T1 IV group relative to the ad libitum-fed 4T1 orthotopic controls but did not differ from the 4T1 orthotopic pair-fed mice (Fig. 2D). Diaphragm, gastrocnemius, tibialis anterior, and soleus masses were likewise reduced in 4T1 IV mice and indistinguishable from 4T1 orthotopic pair-fed animals (Fig. 2D), as were liver, heart, and kidney (Fig. 2E). Lung mass was elevated only in 4T1 IV mice, consistent with their pulmonary tumor burden, whereas splenomegaly was confined to ad libitum-fed 4T1 orthotopic mice (Fig. 2E). The principal exception was brown adipose tissue (BAT), which was slightly but significantly lower in the 4T1 IV group than in either of the 4T1 orthotopic groups (Fig. 2D), indicating that the small BAT mass loss was not accounted for by reduced food intake. Together, these data indicate that anorexia is the predominant driver for tissue wasting in the 4T1 metastasis-induced cancer cachexia model.

### 3.5 Reduced food intake is responsible for muscle atrophy gene induction in the 4T1 model of metastasis-induced cancer cachexia

The ubiquitin ligases *Trim63* (MuRF1) and *Fbxo32* (atrogin-1) and the autophagy-related genes *Bnip3* and *Map1lc3b* are often used as gene markers for muscle atrophy in cancer cachexia^21–23^. We thus examined their expression in our model and its controls. In gastrocnemius, expression of these genes was induced in 4T1 IV mice relative to ad libitum-fed 4T1 orthotopic mice (Fig. 2F). The 4T1 orthotopic pair-fed group, however, showed comparable induction of all four genes, suggesting that their induction in the muscle was driven by reduced food intake. The diaphragm showed the same pattern for *Trim63*, *Fbxo32*, and *Bnip3* across the three groups (Fig. S2A). *Map1lc3b* was the exception: its induction in the diaphragm was specific to the 4T1 IV group. In the heart, expression of these genes was largely unchanged, with only *Bnip3* and *Map1lc3b* modestly elevated in the 4T1 IV mice (Fig. S2B). Altogether, these results indicate that, for elevated expression of the atrophy genes in the 4T1 metastatic model, anorexia is the main contributor.

### 3.5 Muscle weakness is independent of reduced food intake in the model of 4T1 metastasis-induced cancer cachexia

Because loss of lean mass and muscle mass was fully reproduced by pair-feeding (Fig. 2C and D), we next asked whether muscle function was similarly explained by reduced food intake. Forelimb grip strength was significantly reduced in 4T1 IV mice relative to ad libitum-fed orthotopic mice, whereas pair-fed mice retained grip strength (Fig. 3A). Because 4T1 orthotopic pair-fed mice and 4T1 IV mice had lost similar amounts of lean and individual muscle mass (Fig. 2C and D), this deficit in grip strength cannot be attributed to muscle wasting itself, nor was it explained by muscle disuse, as locomotor activity in 4T1 IV mice did not differ from ad libitum-fed orthotopic controls (Fig. 3B and C). Muscle weakness in this model therefore reflects a functional impairment independent of both caloric intake and muscle mass.

**Figure 3.**
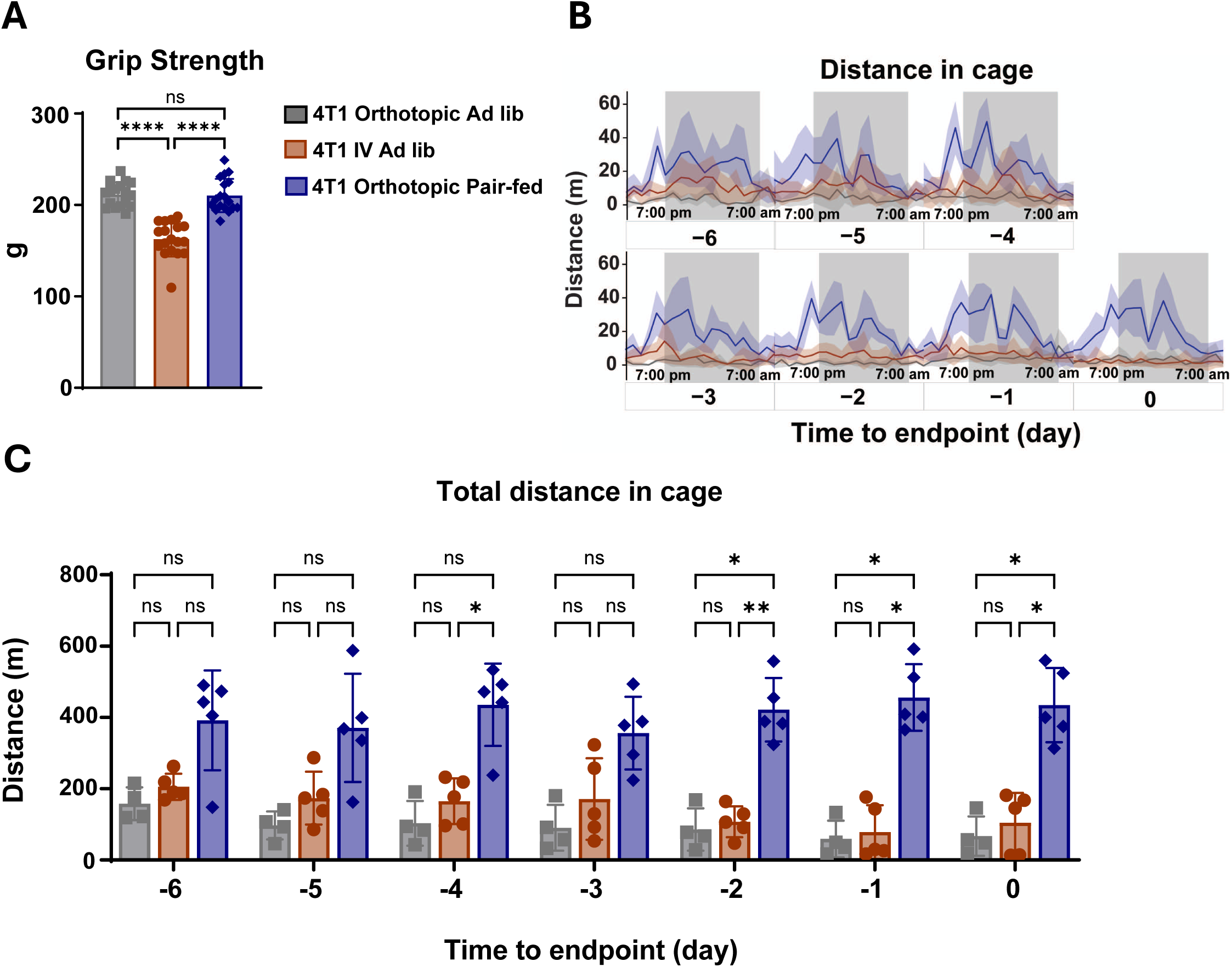
Reduced food intake is not responsible for the reduced muscle function in 4T1 metastasis-induced cancer cachexia. (A) Grip strength in 4T1 orthotopic ad libitum mice (n=17), 4T1 IV ad libitum mice (n=18), and 4T1 orthotopic pair-fed mice (n=18). (B) Home-cage locomotor distance tracked continuously over the final 7 days before endpoint (day-6 to day 0 relative to endpoint), shown across light/dark cycles for each of the three groups: 4T1 orthotopic ad libitum-fed (n=4), 4T1 IV ad libitum-fed (n=4), and 4T1 orthotopic pair-fed groups (n=4). Solid lines represent group means and shaded bands represent SD. (C) Total home-cage distance traveled per day (day-6 to day 0 before endpoint), obtained from the same data in (B). Data are presented as individual data points with mean ± SD in the bar plots. Statistical significance was determined by one-way ANOVA with Tukey’s multiple comparisons. *p□<□0.05; **p□<□0.01; ***p□<□0.001; **** p<0.0001; ns, not significant.

### 3.7 Lower energy expenditure in the 4T1 metastasis-induced model of cancer cachexia is due to reduced food intake

To determine how metastasis alters energy balance in our model, mice were housed in metabolic cages for indirect calorimetry after tumor inoculation. Energy expenditure in 4T1 IV mice started trending down 4 days before the end point (Fig. 4A and B). In the 4T1 orthotopic pair-fed group, energy expenditure showed increasing amplitude of diurnal oscillation approaching the end point, likely due to changing feeding behavior arising from pair-feeding (Fig. 4A). When comparing the daily total of energy expenditure, however, energy expenditure did not differ significantly between the 4T1 IV group and the 4T1 orthotopic pair-fed mice at any timepoint (Fig. 4B), supporting that the decreasing energy expenditure in the 4T1 IV group was due to reduced caloric intake.

**Figure 4.**
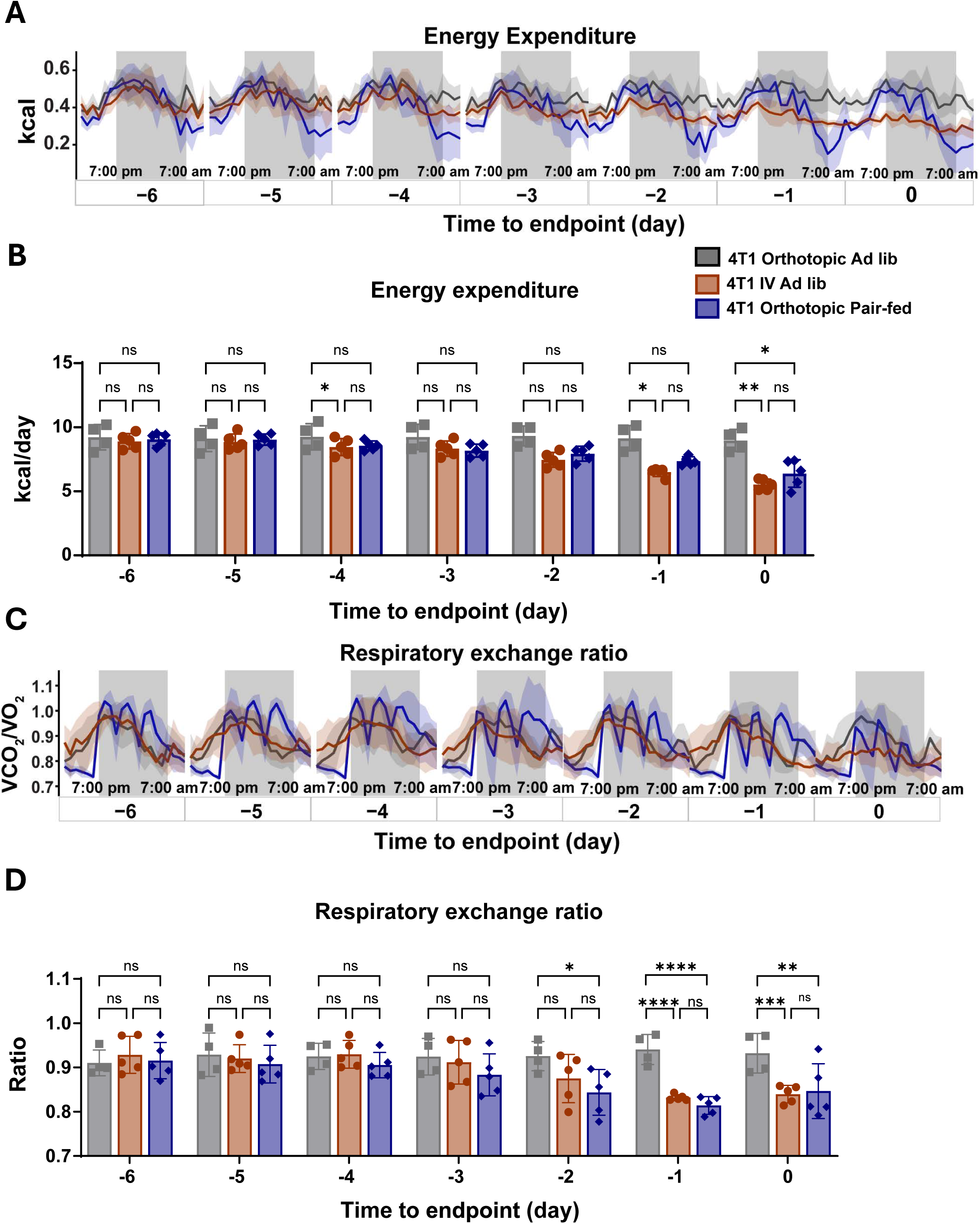
Decline in energy expenditure and respiratory exchange ratio in 4T1 metastasis-induced cancer cachexia is independent of food intake. (A) Energy expenditure measured continuously over the final 7 days before endpoint across light/dark cycles in the three groups: 4T1 orthotopic ad libitum-fed (n=4), 4T1 IV ad libitum-fed (n=4), and 4T1 orthotopic pair-fed groups (n=4). Solid lines represent group means and shaded bands represent SD. (B) Daily energy expenditure for each group across the final 7 days before endpoint. (C) Respiratory exchange ratio (RER, VCO□/VO□) over the same period for these mice for the same mice. Solid lines represent group means and shaded bands represent SD. (D) Averaged daily RER for each group. Data are presented as individual data points with mean ± SD in the bar plots. Statistical significance was determined by one-way ANOVA with Tukey’s multiple comparisons. *p□<□0.05; **p□<□0.01; ***p□<□0.001; **** p<0.0001; ns, not significant.

Respiratory exchange ratio (RER) followed the same trajectory. It started a progressive decline 3-4 days before the end point in both the 4T1 IV group and the 4T1 orthotopic pair-fed group, reaching approximately 0.83 in both groups at the end point (Fig. 4C and D). This decline indicates a shift from carbohydrate toward fat oxidation in the two groups, consistent with negative energy balance. As with energy expenditure, average daily RER did not differ between the 4T1 IV group and the 4T1 orthotopic pair-fed group, demonstrating that the shift in substrate utilization was a consequence of reduced caloric intake.

### 3.8 Elevated levels of circulating IL-6 and TNF-α and impaired metabolic hormone response to reduced food intake in the model of 4T1 metastasis-induced cancer cachexia

To expand our characterization of the model, we profiled serum cytokines and metabolic hormones, which are often implicated as potential mediators of cancer cachexia ^24,25^. Relative to ad libitum-fed 4T1 orthotopic controls, six cytokines were significantly altered in 4T1 IV mice: IL-6, TNF-α, MCP-2, and sICAM-1 were elevated, whereas IL-5 and sCD137/4-1BB/TNFRSF9 were reduced (Fig. 5A, left; Fig. 5B). Comparison against 4T1 orthotopic pair-fed mice then resolved which of these reflected reduced food intake. IL-6 and TNF-α remained significantly elevated relative to pair-fed mice (Fig. 5A, right), identifying them as food intake-independent. IL-5, by contrast, was suppressed at least as strongly in pair-fed mice as in 4T1 IV mice, indicating that its reduction is a consequence of caloric restriction rather than of metastatic disease. For MCP-2, sICAM-1, and sCD137/4-1BB/TNFRSF9, their levels in the 4T1 orthotopic pair-fed group were intermediate and did not differ significantly from either of the other two groups, leaving their food intake dependence unresolved. Thus, we have identified the changes in IL-6 and TNF-α levels as metastasis-specific and food intake-independent features, making them valuable candidates as contributors of cachexia in this model.

**Figure 5.**
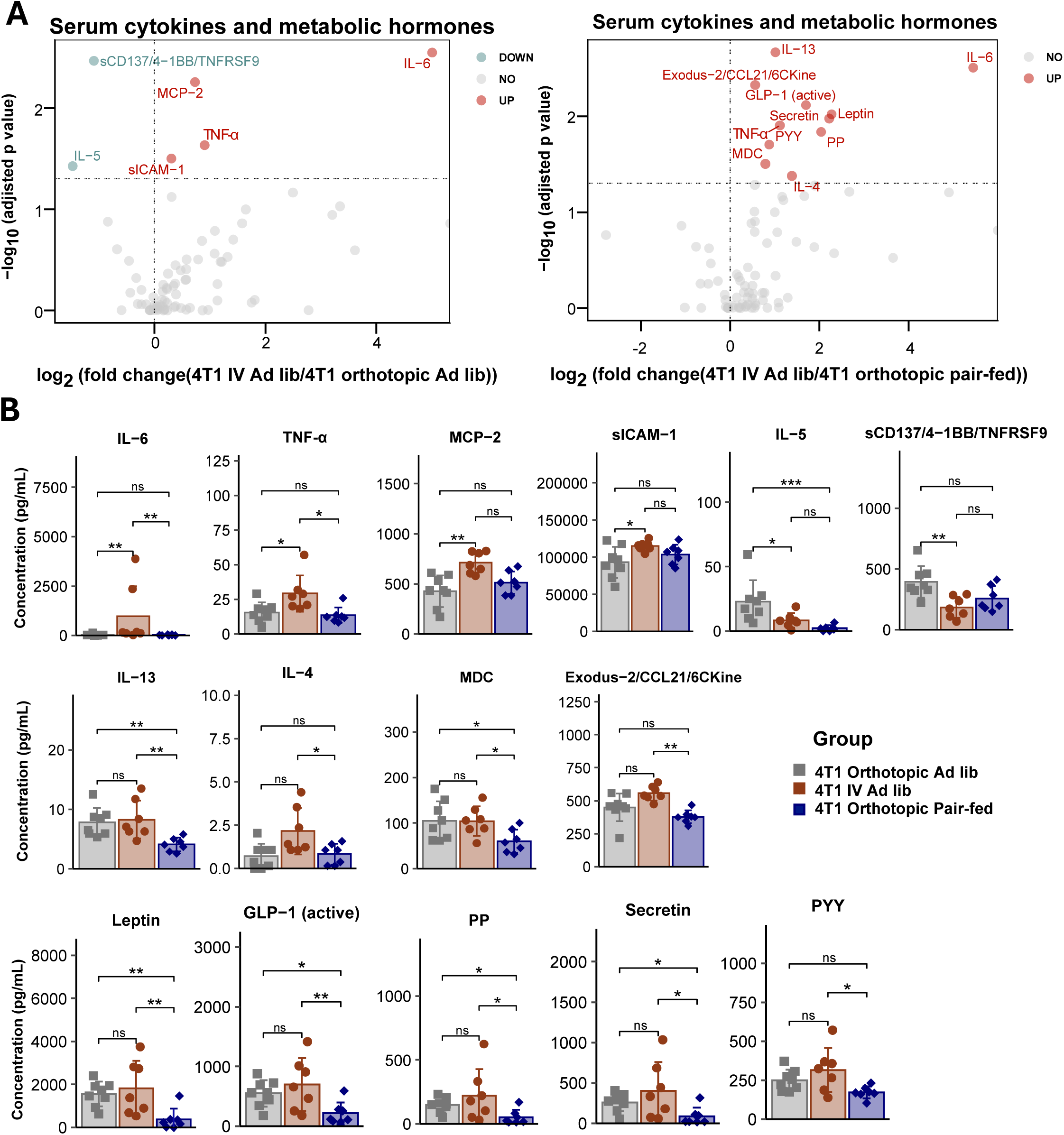
Serum cytokine and metabolic hormone profiling reveals a distinct circulating signature in 4T1 metastasis-induced cancer cachexia. (A) Volcano plot of serum cytokines and metabolic hormones comparing 4T1 IV ad libitum mice vs. 4T1 orthotopic ad libitum mice (left panel), and 4T1 IV ad libitum mice vs. 4T1 orthotopic pair-fed mice (right panel). Analytes are classified as up-, down-, or not significantly regulated. (B) Concentrations of individual analytes that showed significant differences in either comparison. Data are presented as individual data points with mean ± SD in the bar plots. Statistical significance was determined by one-way ANOVA with Tukey’s multiple comparisons. *p□<□0.05; **p□<□0.01; ***p□<□0.001; **** p<0.0001; ns, not significant.

A distinct pattern emerged among analytes that did not differ between 4T1 IV mice and ad libitum-fed 4T1 orthotopic mice. Nine analytes, including IL-13, IL-4, MDC, Exodus-2/CCL21/6CKine, leptin, GLP-1, PP, secretin, and PYY, were significantly lower in pair-fed mice than in 4T1 IV mice (Fig. 5A, right; Fig. 5B). Because the two groups consumed identical amounts of food and were fasted identically before sacrifice, this indicates that 4T1 IV mice lost the endocrine and cytokine suppression that accompanies caloric restriction. Leptin is the clearest example: 4T1 orthotopic pair-fed mice and 4T1 IV mice lost comparable fat mass (Fig. 2C and D), yet leptin was suppressed only in the pair-fed group. Thus, our data revealed that the hormone response to reduced caloric intake is impaired in the model of 4T1 metastasis-induced cancer cachexia, supporting further investigations of its cachectic role.

## 4 Discussion

With this work, we presented a rigorously established and comprehensively characterized model of metastasis-induced cancer cachexia, which we hope for wide adoption by the field for elucidating the mechanistic links from metastatic tumors to cachexia. Although mice bearing 4T1 metastasis have been used to study cachexia, whether wasting is driven by the primary tumor or by metastatic tumor, and whether metastasis acts through the tumor itself or through mechanical compromise of the lung, remain unresolved. We addressed these issues using two isogenic cell lines of differing metastatic capacity delivered either orthotopically or intravenously. Mice bearing orthotopic 4T1 tumors maintained body weight, whereas mice with established 4T1 lung metastases developed severe cachexia; 67NR cells, which colonize the lung efficiently when delivered intravenously, produced no wasting at pulmonary burdens comparable to those of the 4T1 model. Cachexia in this model therefore requires 4T1 cells growing in the lung and is attributable neither to 4T1 tumor burden itself nor to the presence of tumor mass in the lung, unambiguously establishing that cachexia in this model is caused by 4T1 metastatic tumors.

In addition to a robust model of metastasis-induced cancer cachexia, our work further established a strict control group, mice bearing 4T1 tumors at the orthotopic primary site. As it is the same cancer cells in primary tumors and metastatic tumors, comparing measurements between cachectic mice to this control will reveal changes that are specific to metastasis and cachexia. Without such a control, the list of changes will be much longer, including any changes that arise from 4T1 cancer cells. Moreover, we demonstrated that pair-feeding mice bearing the 4T1 primary tumors additionally controls for effects from reduced food intake, further narrowing down the list of measured changes in the cachectic group. Therefore, with the 4T1 metastatic group and the control group of food intake-matched 4T1 primary tumor-bearing mice, we provide a strong platform for identifying metastasis-specific cachectic factors and pathways.

As with all preclinical models, the model here has limitations. One is the IV injection approach to induce metastasis. Intravenous injection models lung colonization rather than the complete metastatic cascade, bypassing invasion and intravasation, so the phenotypes we describe are attributable to established metastases rather than to the process of metastasizing. Another limitation is the rapid development of cancer and cachexia, which may not capture important elements in the much slower process associated with spontaneous cancer. Finally, we used tumor mass as a measure of tumor burden

## 5 Conclusion

Through systematic comparison of primary and metastatic tumor models using isogenic 4T1 and 67NR cell lines, we established a cancer cachexia model in which cachexia is specifically associated with 4T1 lung metastasis rather than primary tumor growth or lung tumor burden alone. Comprehensive phenotyping demonstrated that this model recapitulates key features of clinical cachexia, including anorexia, weight loss, loss of adipose and lean mass, skeletal muscle wasting, and reduced grip strength. By incorporating a genetically identical, food intake-matched primary tumor control, we further distinguished the effects of metastatic disease from those secondary to tumor burden or reduced food intake. These studies revealed that anorexia accounts for much of the tissue wasting, whereas muscle weakness and alterations in select circulating factors, including IL-6, TNF-α, and leptin, occur independently of reduced food intake. Together, these findings establish the 4T1 lung metastasis model and its matched primary tumor control as a rigorously characterized platform for identifying metastasis-specific mediators of cachexia and provide a framework for studying cancer cachexia in the metastatic setting in which it most commonly manifests clinically.

## Acknowledgments

This work was delivered as part of the CANCAN Team supported by the Cancer Grand Challenges partnership funded by Cancer Research UK (CGCATF-2021/100034) and the National Cancer Institute (OT2CA278654), and Find the Cause Breast Cancer Foundation (J.M.U.).

## Ethics Statement

All animal care and experimental procedures were approved by the Institutional Animal Care and Use Committees (IACUC) of the Harvard Medical Area (Protocol #: IS00002982-6).

## Data Availability Statement

The data that support the findings of this study are available from the corresponding author upon reasonable request.

---

**Figure S1.**
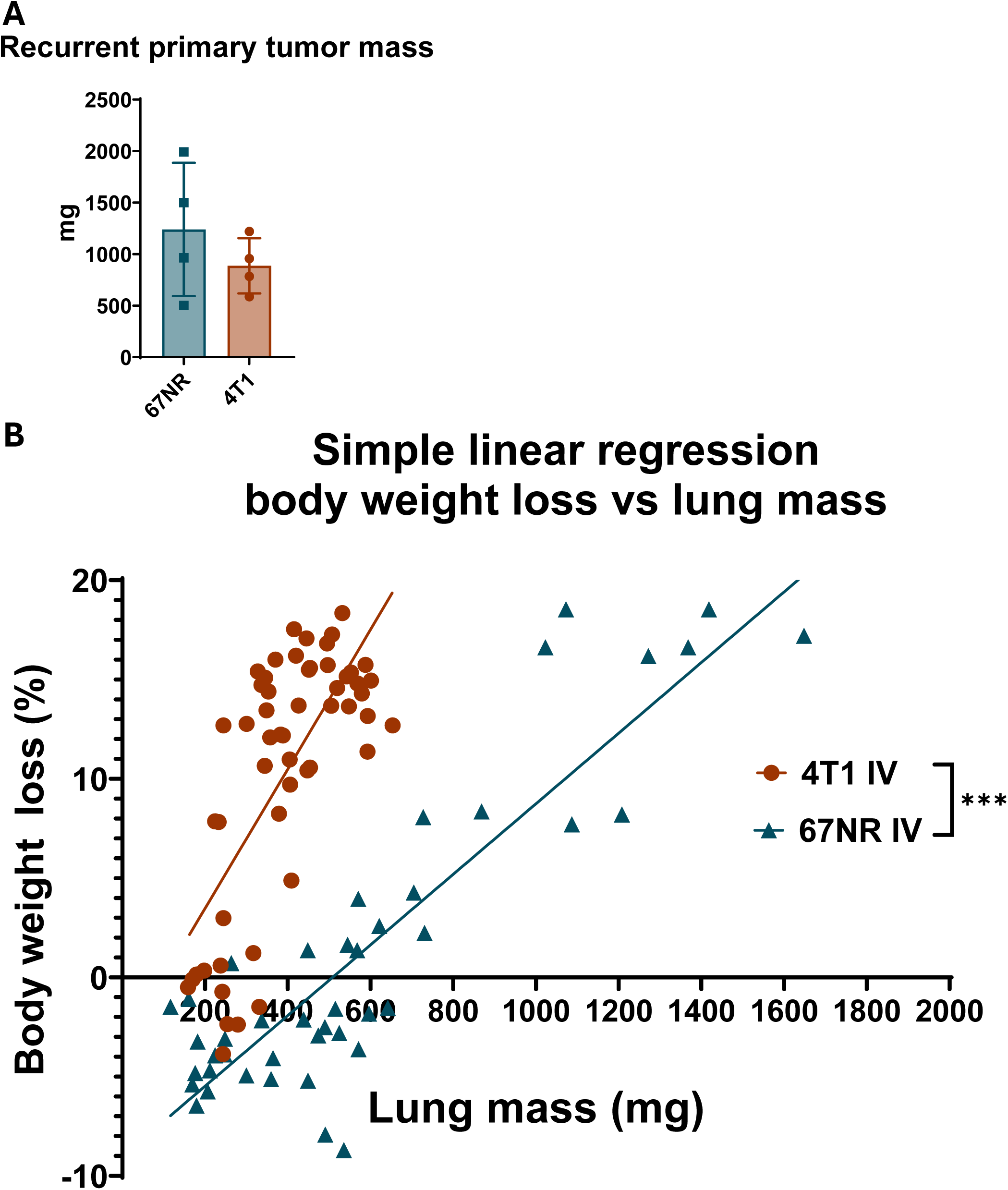
Orthotopic tumor recurrence after surgical resection and relationship between lung metastatic burden and weight loss. (A) Recurrent primary tumor mass at experimental endpoint in mice who underwent surgical resection of the primary orthotopic tumor with 67NR (n=4) or 4T1 (n=4) cells. Primary tumors were resected at day 12 post-tumor inoculation. Data are presented as individual data points with mean ± SD in the bar plot. (B) Linear regression of body weight loss versus lung mass of mice injected intravenously with 4T1 (n=54) or 67NR (n=44) cells, showing lung mass-independent weight loss in the 4T1 IV mice. Statistical significance was determined by simple linear regression analysis. *p□<□0.05; **p□<□0.01; ***p□<□0.001; **** p<0.0001; ns, not significant.

**Figure S2.**
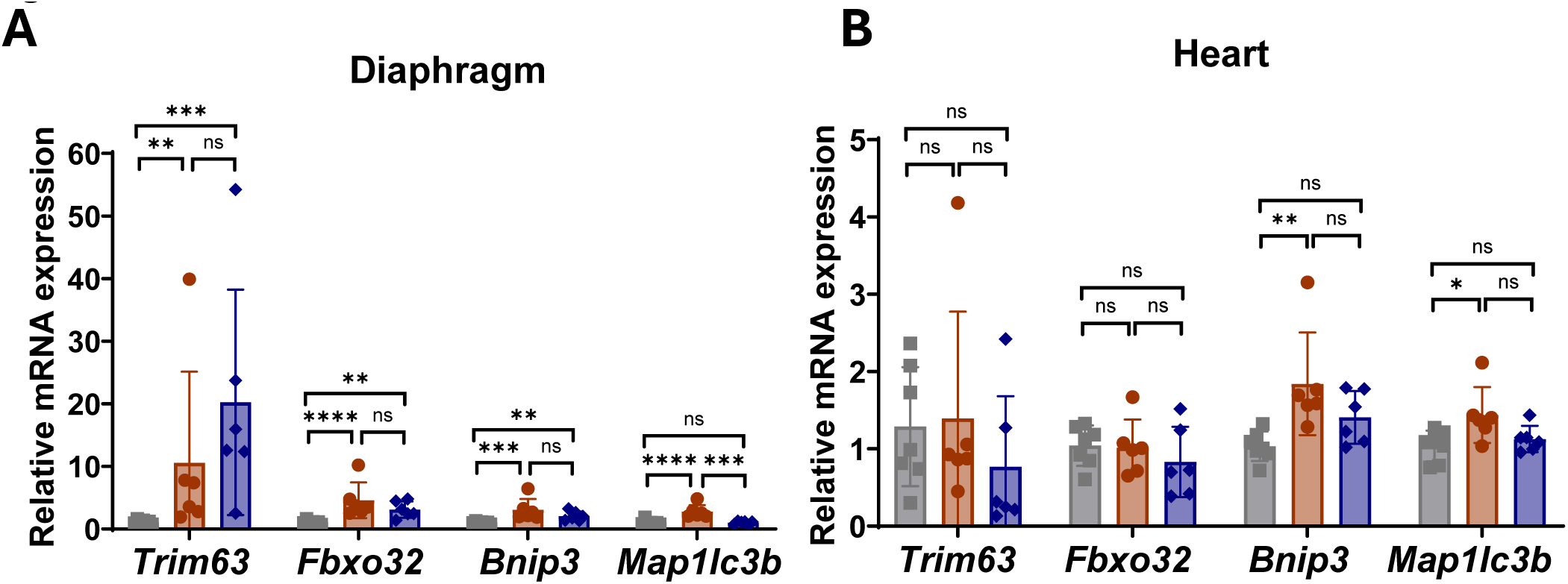
Muscle atrophy and autophagy gene expression in (A) diaphragm and (B) heart from 4T1 orthotopic ad libitum-fed (n=6), 4T1 IV ad libitum-fed (n=7), and 4T1 orthotopic pair-fed groups (n=6). Data are presented as individual data points with mean ± SD in the bar plots. Statistical significance was determined by one-way ANOVA with Tukey’s multiple comparisons. *p□<□0.05; **p□<□0.01; ***p□<□0.001; **** p<0.0001; ns, not significant.

**Table 1.**
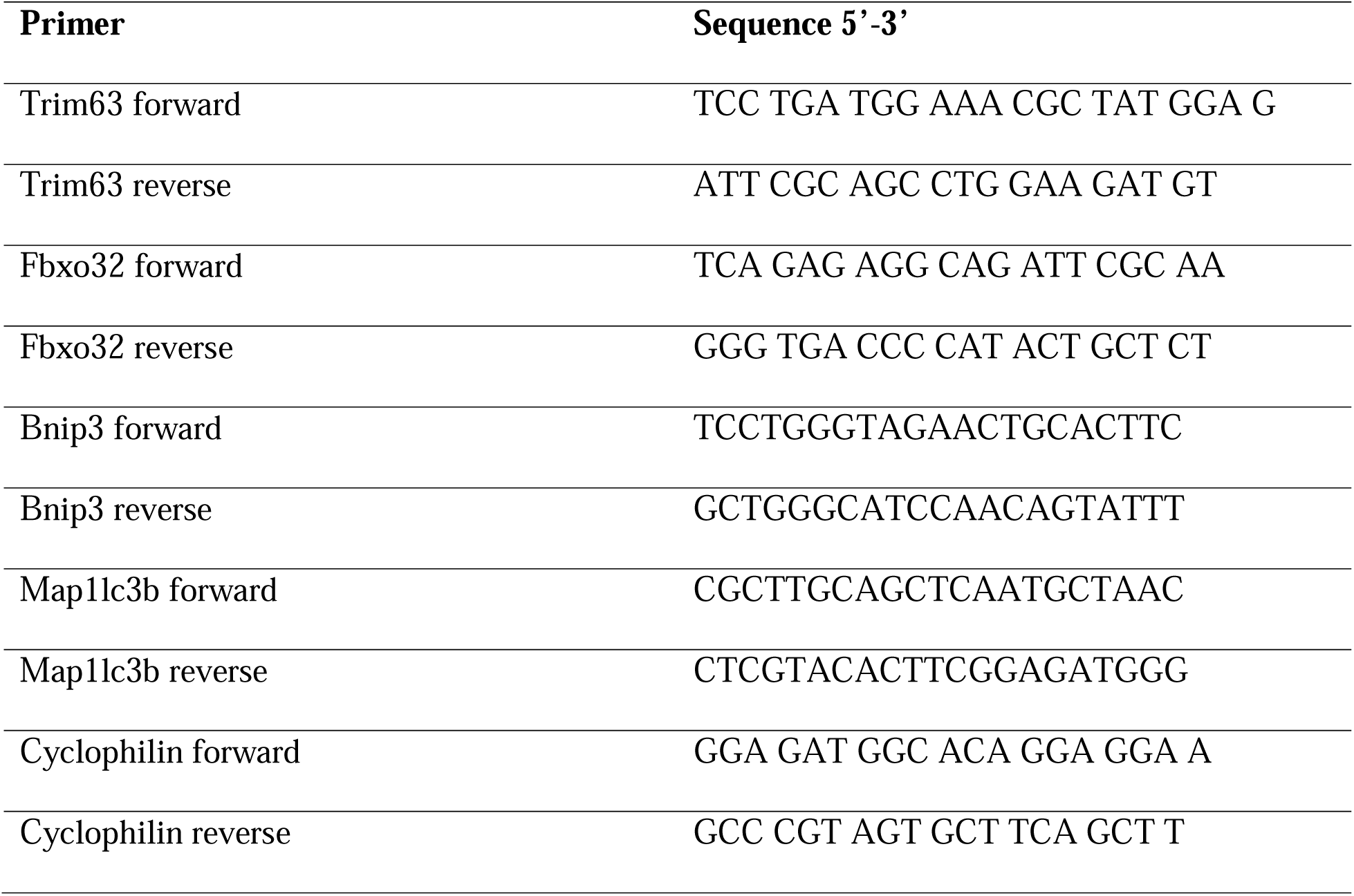
Primer sequences.

